# In Silico Analysis and Modeling of Novel Pathogenic Single Nucleotide Polymorphisms (SNPs) in Human *CD40LG* Gene

**DOI:** 10.1101/552596

**Authors:** Abdelrahman H. Abdelmoneim, Mujahed I. Mustafa, Thwayba A. Mahmoud, Naseem S. Murshed, Mohamed A. Hassan

## Abstract

**Background:** The X-linked hyper-immunoglobulin M syndrome (XHIGM) is a rare, inherited immune deficiency disorder. It is more common in males. Characterized by elevated serum IgM levels and low to undetectable levels of serum IgG, IgA and IgE. Hyper-IgM syndrome is caused by mutations in the *CD40LG* gene. Located in human Xq26. CD40LG acts as an immune modulator in activated T cells.

**Method:** We used different bioinformatics tools to predict the effect of each SNP on the structure and function of the protein.

**Result:** 8 novel SNPs out of 233 were found to have most deleterious effect on the protein structure and function. *While modeling of* nsSNPs was studied by Project HOPE software.

**Conclusion:** Better understanding of Hyper-IgM syndrome caused by mutations in CD40LG gene was achieved using in silico analysis. This is the first in silico functional analysis of CD40LG gene and 8 novel mutations were found using different bioinformatics tools, and they could be used as diagnostic markers for hyper-IgM syndrome. These 8 novel SNPs may be important candidates for the cause of different types of human diseases by *CD40LG* gene.

## Introduction

The X-linked hyper-immunoglobulin M syndrome (XHIGM) is a rare, inherited immune deficiency disorder.[1-6] autosomal recessive HIES (AR-HIES) is the most frequent form of the Hyper IgM syndrome which is characterized by elevated serum IgM levels and low to undetectable levels of serum IgG, IgA and IgE..[7-10] it is more common in males. [10-15]

Hyper-IgM syndrome is caused by mutations in the *CD40LG* gene.[16-19] located in human Xq26.[20] *CD40LG* acts as an immune modulator in activated T cells[21] Inactivation of *CD40LG* gene resulted from the insertion of an AluYb8 element in exon 1 responsible for a total deficiency of CD40 ligand expression by T lymphocytes[22] Interestingly, some study shows that, not all mutations occur in *CD40LG* gene may cause XHIGM [23] Different mutations have been reported. [23-32]

As far as we know there are two options for treatment, either hematopoietic stem cell transplantation or intravenous immunoglobulin replacement therapy. [33] but it is sometimes completely ignored because of low recognition and limited knowledge of this rare disease.[34] Mutations in *CD40LG* gene may also cause: Sickle cell anemia, systemic lupus erythematosus, malignancies, and discusses neuroendocrine tumors (NETs) arising in other immunocompromised states. Of all primary immune deficiency diseases, NETs appear to be distinctive to XHIGM patients. An outcome for XHIGM patients with NETs is poor, and the mechanism behind this association remains unknown.[35-37]

The aim of this study is to identify functional SNPs within dbSNP located in coding region of *CD40LG* gene using in silico analysis. The usage of in silico analysis has strong influence on the identification of candidate SNPs since they are easy and less costly, and can assist in pharmacogenomics by identifying high risk SNP variants contributing to drug response as well as developing novel therapeutic elements and diagnostic markers. [38-42]

## Materials and Methods

### Data mining

The data on human *CD40LG* gene was collected from National Center for Biological Information (NCBI) web site.(https://www.ncbi.nlm.nih.gov/) and the protein sequence was collected from Uniprot (https://www.uniprot.org/).

### SIFT

We used SIFT to observe the effect of A.A. substitution on protein function. SIFT predicts damaging SNPs on the basis of the degree of conserved amino A.A. residues in aligned sequences to the closely related sequences, gathered through PSI-BLAST.[43] It is available at (http://sift.jcvi.org/).

### PolyPhen

PolyPhen (version 2) We used PolyPhen to study probable impacts of A.A. substitution on structural and functional properties of the protein by considering physical and comparative approaches.[44] It is available at (http://genetics.bwh.harvard.edu/pph2/).

### Provean

Provean is a software tool which predicts whether an amino acid substitution or indel has an impact on the biological function of a protein. It is useful for filtering sequence variants to identify nonsynonymous or indel variants that are predicted to be functionally important.[45] It is available at (https://rostlab.org/services/snap2web/).

### SNAP2

SNAP2 is a trained classifier that is based on a machine learning device called “neural network”. It distinguishes between effect and neutral variants/non-synonymous SNPs by taking a variety of sequence and variant features into account.[46] It is available at (https://rostlab.org/services/snap2web/).

### SNPs&GO

SNPs&GO is an accurate method that, starting from a protein sequence, can predict whether a variation is disease related or not by exploiting the corresponding protein functional annotation. SNPs&GO collects in unique framework information derived from protein sequence, evolutionary information, and function as encoded in the Gene Ontology terms, and outperforms other available predictive methods.[47] It is available at (http://snps.biofold.org/snps-and-go/snps-and-go.html).

### PHD-SNP

An online Support Vector Machine (SVM) based classifier, is optimized to predict if a given single point protein mutation can be classified as disease-related or as a neutral polymorphism, it is available at: (http://snps.biofold.org/phd-snp/phdsnp.html).

### I-Mutant 3.0

I-Mutant 3.0 Is a neural network based tool for the routine analysis of protein stability and alterations by taking into account the single-site mutations. The FASTA sequence of protein retrieved from UniProt is used as an input to predict the mutational effect on protein stability. [48] It is available at (http://gpcr2.biocomp.unibo.it/cgi/predictors/I-Mutant3.0/I-Mutant3.0.cgi).

### MUpro

MUpro is a support vector machine-based tool for the prediction of protein stability changes upon nonsynonymous SNPs. The value of the energy change is predicted, and a confidence score between −1 and 1 for measuring the confidence of the prediction is calculated. A score <0 means the variant decreases the protein stability; conversely, a score >0 means the variant increases the protein stability.[49] It is available at (http://mupro.proteomics.ics.uci.edu/).

### GeneMANIA

We submitted genes and selected from a list of data sets that they wish to query. GeneMANIA approach to know protein function prediction integrate multiple genomics and proteomics data sources to make inferences about the function of unknown proteins.[50] It is available at (http://www.genemania.org/).

### Structural Analysis

#### Developing 3D structure of mutant STAT3 gene

The 3D structure of human CD40LG protein is not available in the Protein Data Bank. Hence, we used RaptorX to generate a 3D structural model for wild-type CD40LG. RaptorX is a web server predicting structure property of a protein sequence without using any templates. It outperforms other servers, especially for proteins without close homologs in PDB or with very sparse sequence profile.[51] It is available at (http://raptorx.uchicago.edu/).

#### Modeling Amino Acid Substitution

UCSF Chimera is a highly extensible program for interactive visualization and analysis of molecular structures and related data, including density maps, supramolecular assemblies, sequence alignments, docking results, conformational analysis Chimera (version 1.8).[52] It is available at (http://www.cgl.ucsf.edu/chimera/).

#### Identification of Functional SNPs in Conserved Regions by using ConSurf server

ConSurf web server provides evolutionary conservation profiles for proteins of known structure in the PDB. Amino acid sequences similar to each sequence in the PDB were collected and multiply aligned using CSI-BLAST and MAFFT, respectively. The evolutionary conservation of each amino acid position in the alignment was calculated using the Rate 4Site algorithm, implemented in the ConSurf web server. The algorithm takes explicitly into account the phylogenetic relations between the aligned proteins and the stochastic nature of the evolutionary process. Rate 4 Site assigns a conservation level for each residue using an empirical Bayesian inference. Visual inspection of the conservation patterns on the 3-dimensional structure often enables the identification of key residues that comprise the functionally-important regions of the protein.[53, 54] It is available at (http://consurf.tau.ac.il/).

## Results

## Discussion

8 novel mutations were found to have a damaging effect on the stability and function of the *CD40LG* gene using bioinformatics tools. The human immune system, especially the adaptive branch, significantly declines with ageing. Several distinct immunosenescent measures have already been described, yet data regarding to age-associated baseline alterations in immune cell function is limited.[55] Therefore, We investigated the effect of each SNP on the function and stability of the protein and gene expression of immune-related genes using different softwares with different parameters and aspects, in order to confirm the results we found and to minimize the error to the least percentage possible. The software used were SIFT, Polyphen-2, PROVEAN, SNAP2, PhD-SNP, SNP&GO, I-Mutant 3.0, MUPro, and Project HOPE. (figure 1)

**Figure 1:**
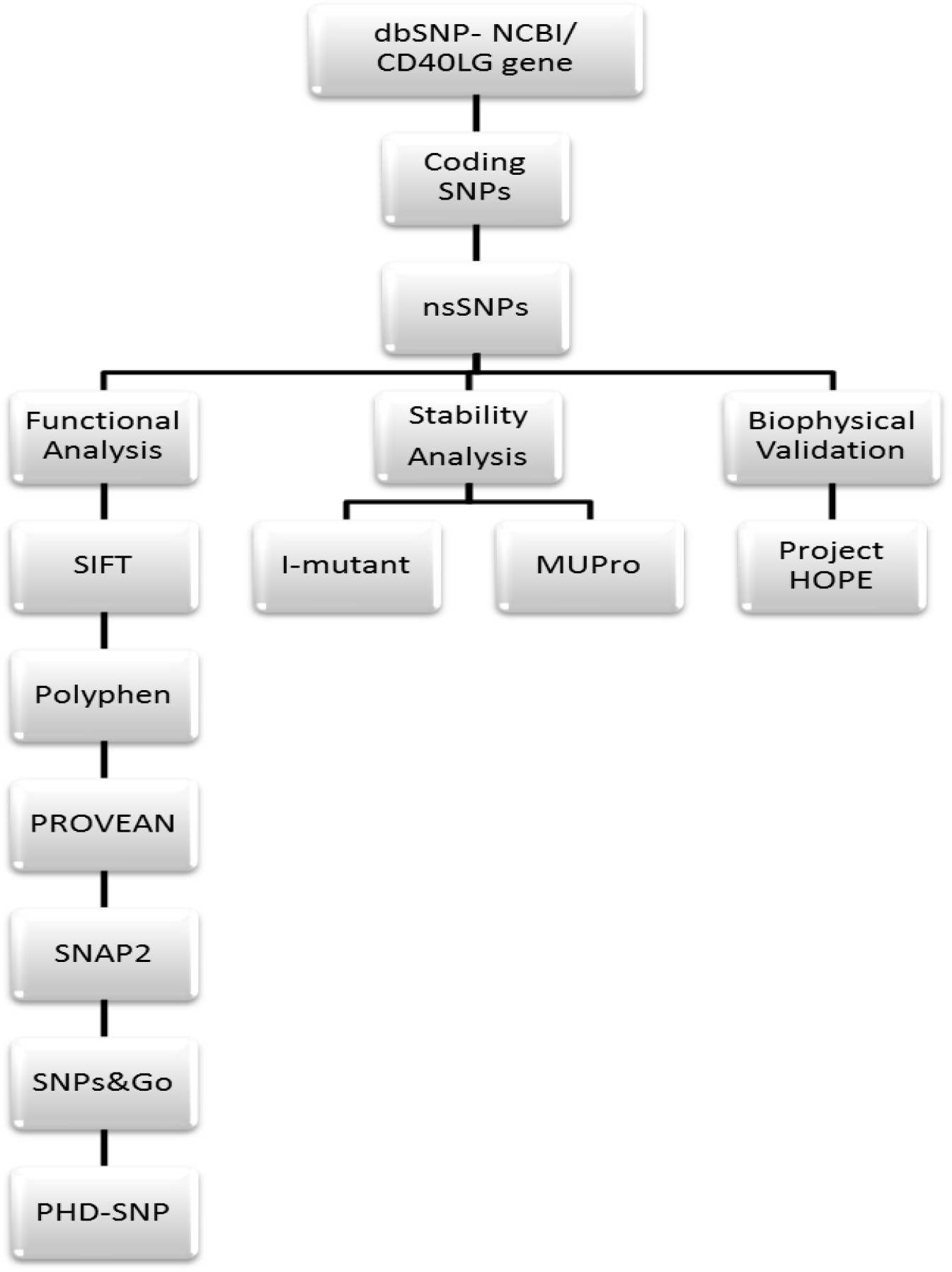
Workflow Software’s used in SNPs analysis.

**Figure 2:**
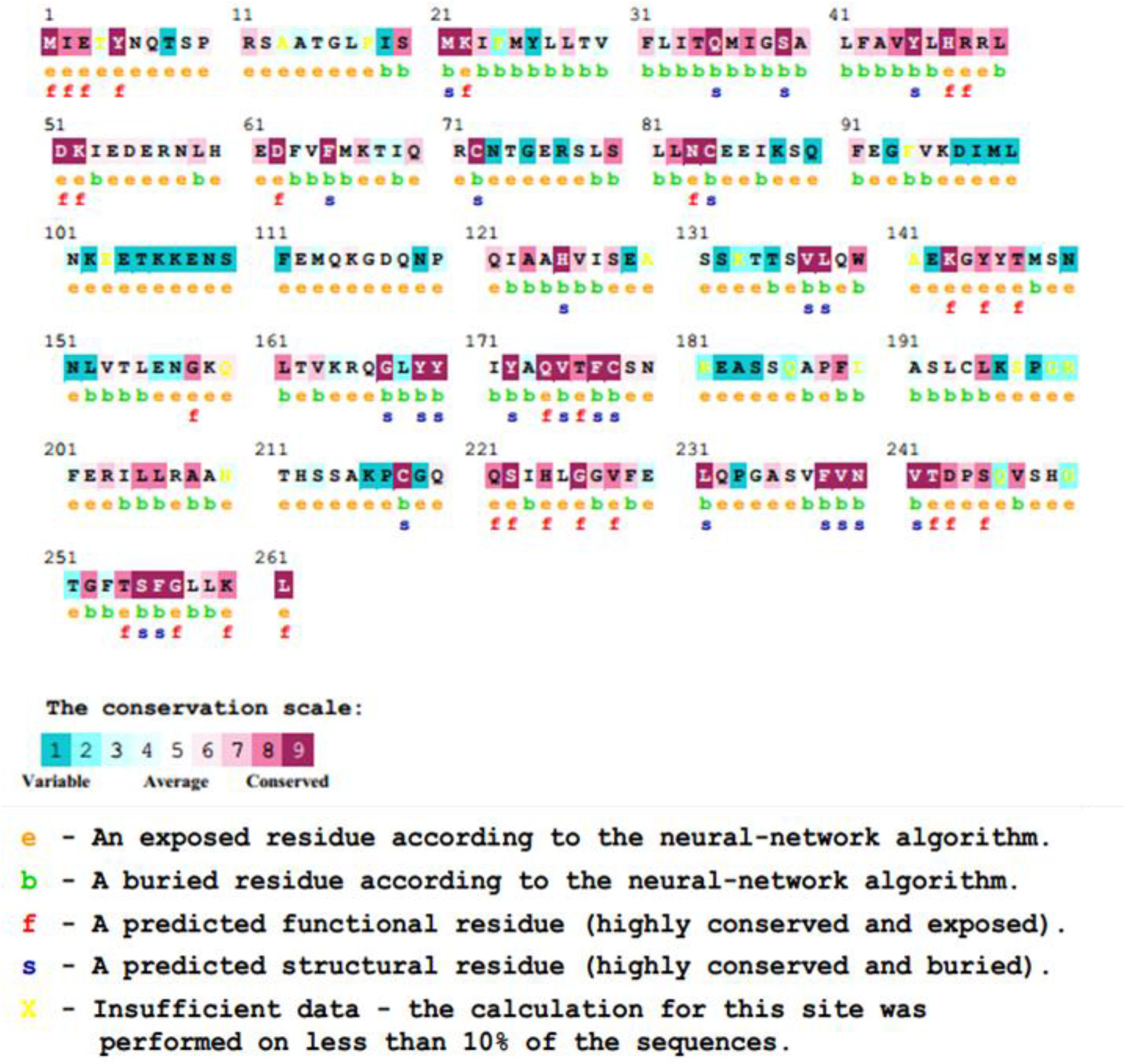
Unique and conserver regions in CD40LG protein were determined using Consurf.

**Figure 3:**
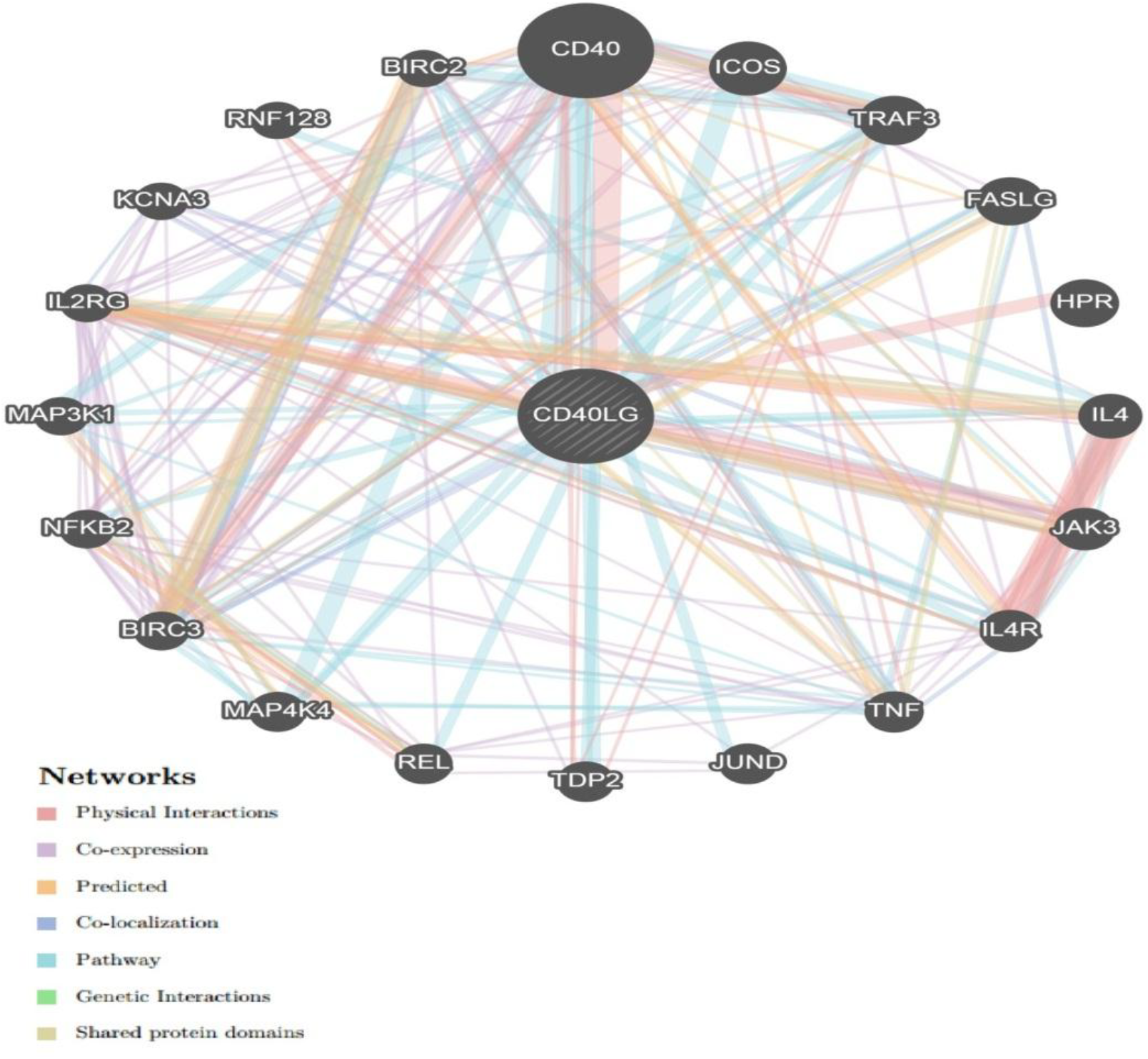
Interaction between *CD40LG* and its related genes.

We retrieved 322 SNPs from the dbSNP/NCBI Database, which was the total number of nsSNPs in the coding region of the *CD40LG* gene in *Homo sapiens*. There were 105 nsSNPs (missense mutations) then submitted them to functional analysis by SIFT, PolyPhen-2, PROVEAN and SNAP2. SIFT server predicted 38 deleterious SNPs, polyphen-2 predicted 56 damaging SNPs (19 possibly damaging and 37 probably damaging), PROVEAN predicted 23 deleterious SNPs and SNAP2 predicted 53 deleterious SNPs. After filtering the Quadra-positive deleterious SNPs we ended up with 15 SNPs (Table 1) and we submitted them to PhD-SNP and SNP&GO to further investigate their effect on the function. PhD-SNP predicted 15 disease-associated SNPs while SNP&GO predicted 8, so we filtered the double positive 8 SNPs (Table 2) and submitted them to I-Mutant 3.0 and MUPro to investigate their effect on the stability. All the SNPs were found to cause a decrease in the stability of the protein except for one SNP (G257D) predicted by I-Mutant 3.0 to increase the stability. (Table 3)

**Table (1):** Damaging or Deleterious nsSNPs associated variations predicted by various softwares:

| dbSNP rs# | SUB | SIFT<br>PREDICTION | SIFT<br>SCORE | POLYPHEN<br>PREDICTION | POLYPHEN<br>SCORE | Prediction<br>(cutoff= -<br>2.5) | PROVEAN<br>score | SNAP2<br>PREDICTION | SNAP2<br>SCORE |
| --- | --- | --- | --- | --- | --- | --- | --- | --- | --- |
| rs750090324 | Y5C | Damaging | 0 | probably<br>damaging | 0.999 | Deleterious | -3.085 | effect | 64 |
| rs104894778 | A123E | Damaging | 0 | probably<br>damaging | 1 | Deleterious | -3.3 | effect | 97 |
| - | A123E | Damaging | 0 | probably<br>damaging | 1 | Deleterious | -3.3 | effect | 97 |
| rs104894777 | W140G | Damaging | 0 | probably<br>damaging | 1 | Deleterious | -9.404 | effect | 96 |
| rs886039326 | G144E | Damaging | 0.02 | probably<br>damaging | 0.993 | Deleterious | -3.333 | effect | 70 |
| rs773698908 | M148K | Damaging | 0.02 | possibly<br>damaging | 0.501 | Deleterious | -3.107 | effect | 85 |
| rs104894769 | L155P | Damaging | 0.01 | probably<br>damaging | 1 | Deleterious | -3.089 | effect | 92 |
| - | L155P | Damaging | 0.01 | probably<br>damaging | 1 | Deleterious | -3.089 | effect | 92 |
| rs786205606 | Y169C | Damaging | 0 | probably<br>damaging | 1 | Deleterious | -8.868 | effect | 60 |
| rs756468554 | Y170C | Damaging | 0 | probably<br>damaging | 1 | Deleterious | -7.592 | effect | 69 |
| rs200840715 | R181W | Damaging | 0 | probably<br>damaging | 0.999 | Deleterious | -2.675 | effect | 65 |
| rs993299705 | I223F | Damaging | 0.01 | probably<br>damaging | 0.998 | Deleterious | -2.768 | effect | 62 |
| rs104894768 | G227V | Damaging | 0 | probably<br>damaging | 1 | Deleterious | -6.598 | effect | 95 |
| rs1313764088 | V237A | Damaging | 0.01 | probably<br>damaging | 0.968 | Deleterious | -3.208 | effect | 40 |
| rs193922136 | T254M | Damaging | 0.01 | probably<br>damaging | 1 | Deleterious | -4.261 | effect | 66 |
| rs1057521128 | F256S | Damaging | 0.01 | probably<br>damaging | 1 | Deleterious | -7.425 | effect | 86 |
| rs1477466218 | G257D | Damaging | 0 | probably<br>damaging | 1 | Deleterious | -6.453 | effect | 86 |

**Table (2):**
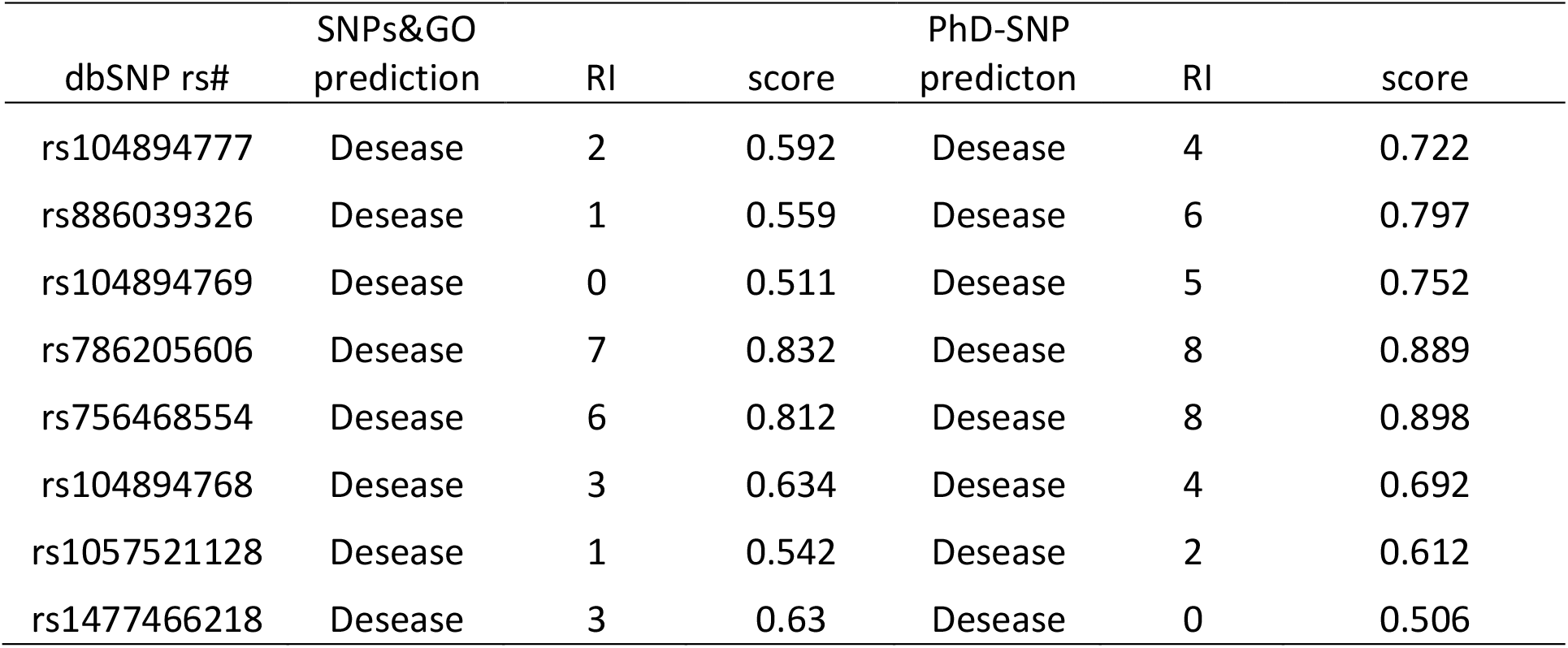
Disease effect nsSNPs associated variations predicted by SNPs&GO and PhD-SNP:

**Table (3):** stability analysis predicted by I-Mutant version 3.0 and MUPro (also Show the 8 novel mutations):

| dbSNP rs# | SUB | SVM2 Prediction Effect | RI | DDG Value Prediction | Mupro Prediction |
| --- | --- | --- | --- | --- | --- |
| rs104894777 | W140G | Decrease | 9 | -2.06 | Decrease |
| rs886039326 | G144E | Decrease | 1 | -0.68 | Decrease |
| rs104894769 | L155P | Decrease | 6 | -1.6 | Decrease |
| rs786205606 | Y169C | Decrease | 2 | -1.14 | Decrease |
| rs756468554 | Y170C | Decrease | 3 | -1.3 | Decrease |
| rs104894768 | G227V | Decrease | 1 | -0.14 | Decrease |
| rs1057521128 | F256S | Decrease | 6 | -1.5 | Decrease |
| rs1477466218 | G257D | Increase | 0 | -0.69 | Decrease |

We submitted the most deleterious 8 SNPs to project HOPE which revealed that all the SNPs are located in a domain in the protein and thus might have a vital change in the structure and function of the protein and it may affect its ability to bind with its targets. And we used Chimera software to visualize the amino acids change (figures 4-11)

**Figure 4:**
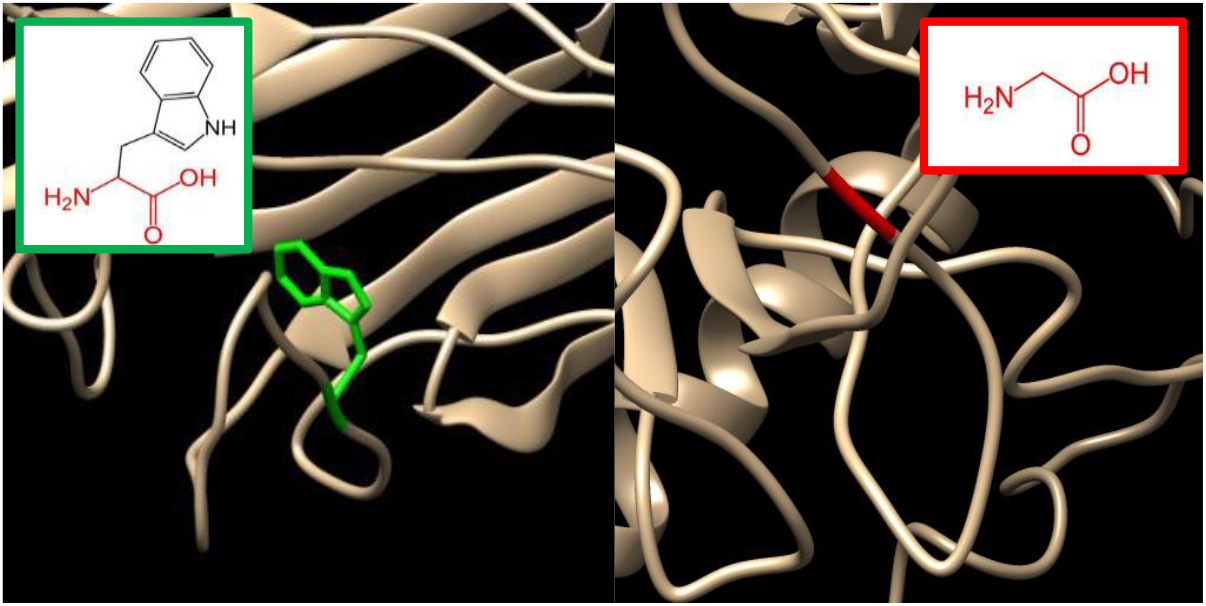
(W140G): The amino acid Tryptophan change to Glycine at position 140.

**Figure 5:**
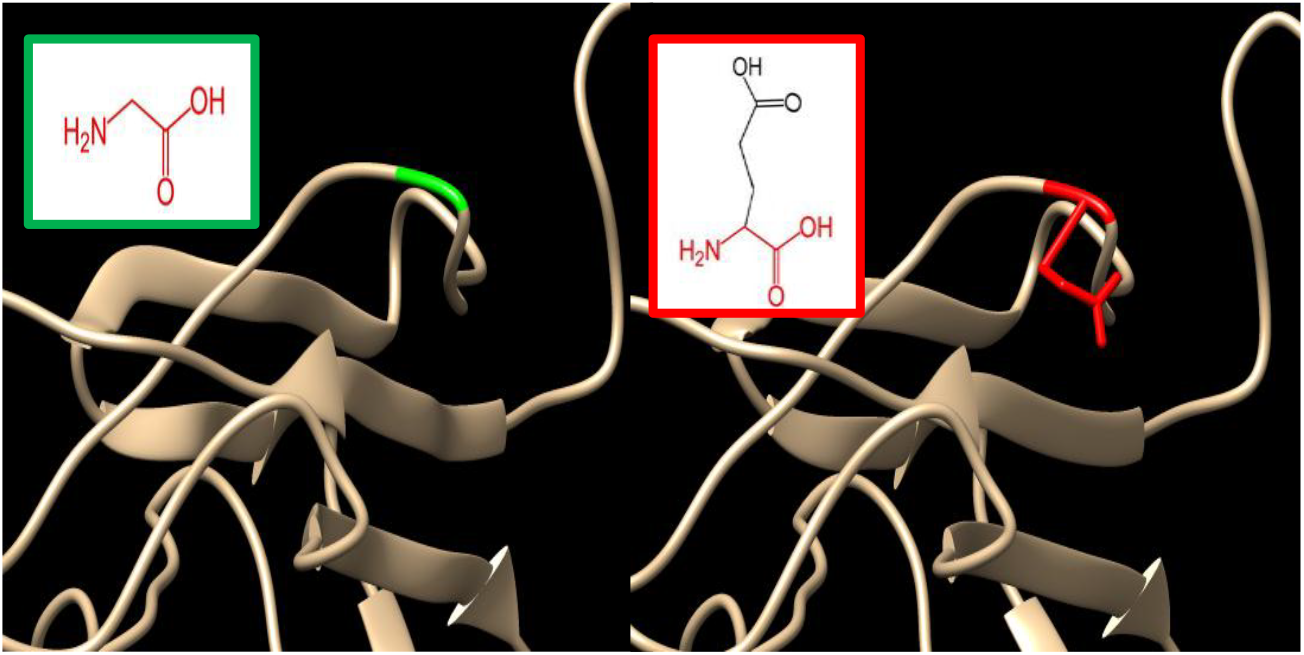
(G144E) The amino acid Glycine change to Glutamic Acid at position 144.

**Figure 6:**
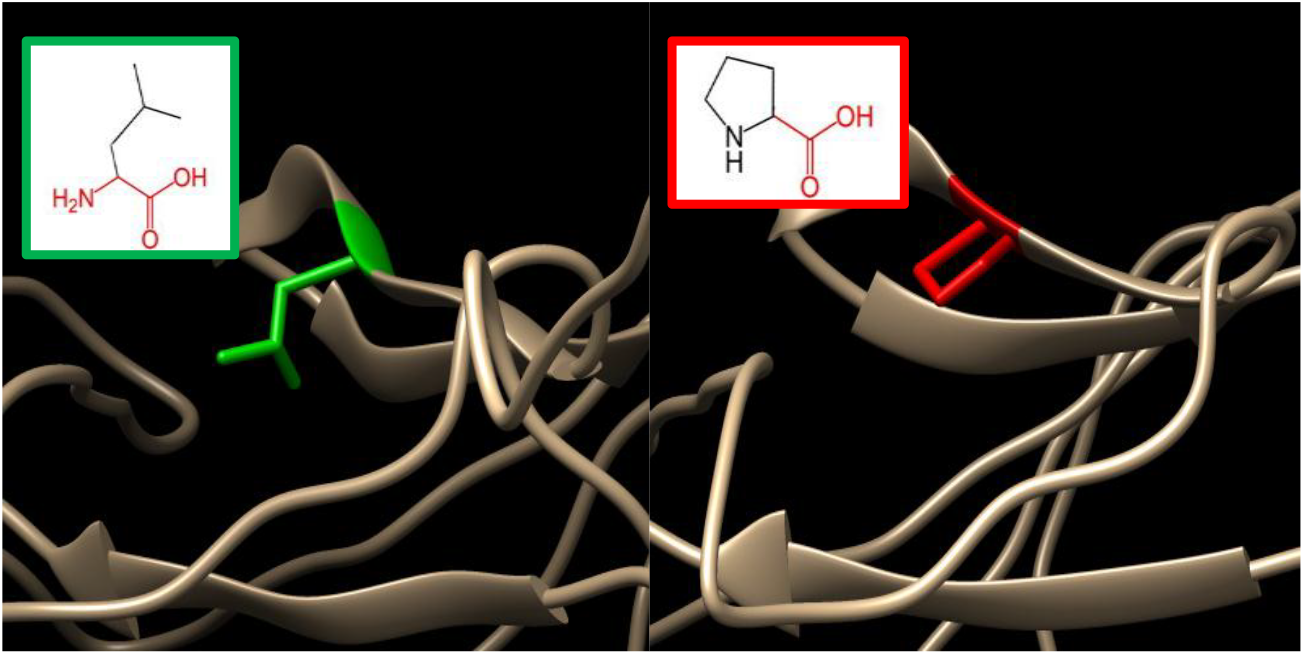
(L155P)The amino acid Leucine change to Proline at position 155.

**Figure 7:**
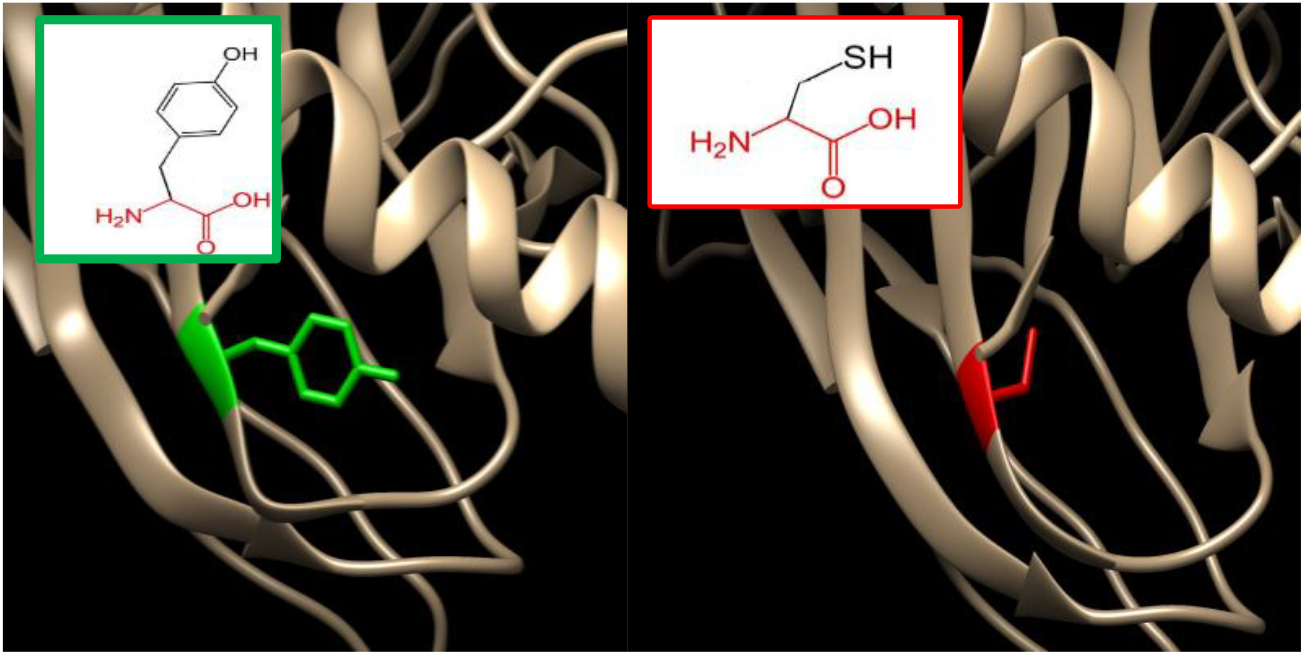
(Y169C)The amino acid Tyrosine change to Cysteine at 169.

**Figure 8:**
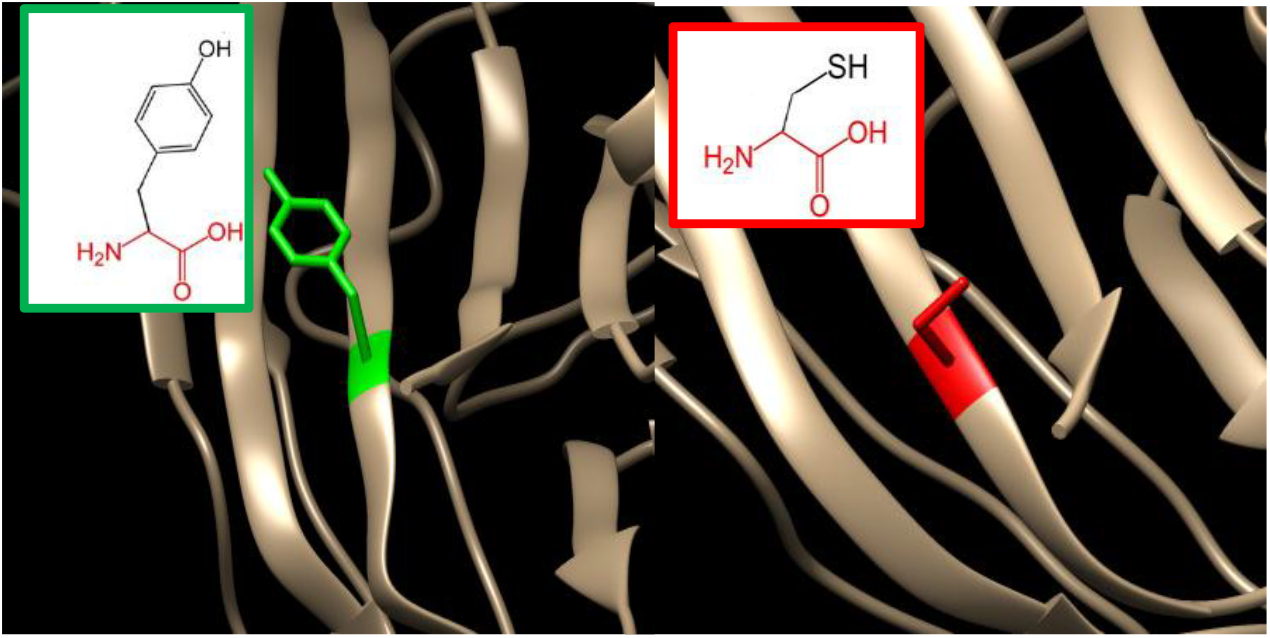
(Y170C) The amino acid Tyrosine change to Cysteine at 170.

**Figure 9:**
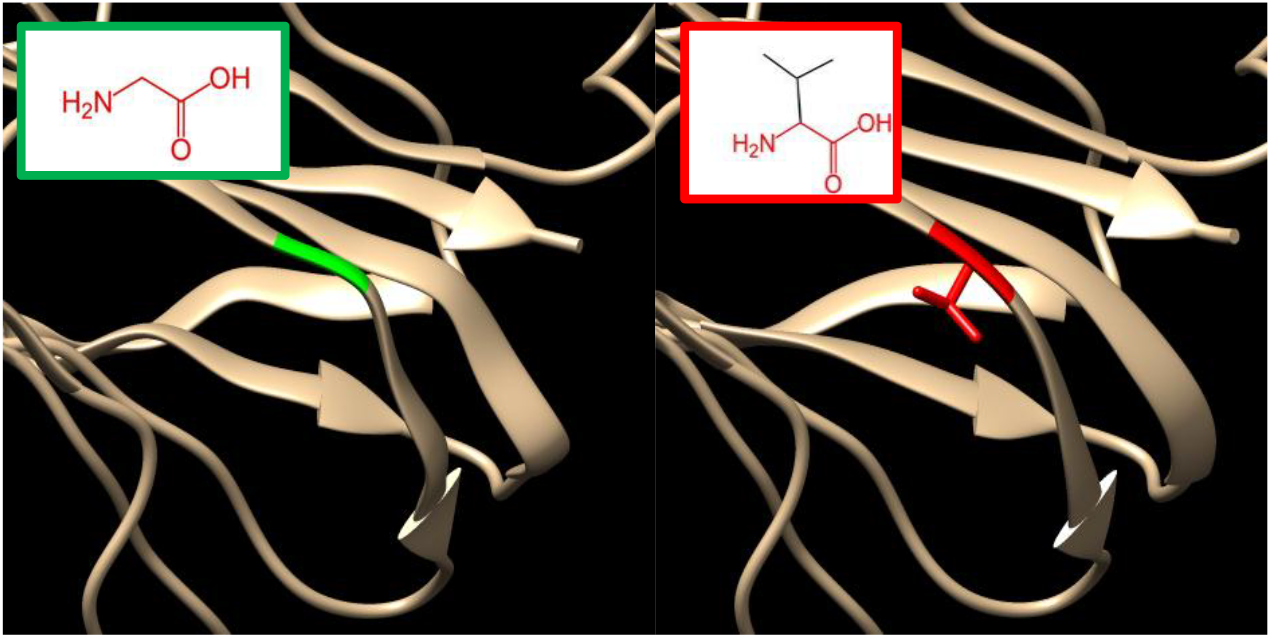
(G227V) The amino acid Glycine change to Valine at 227.

**Figure 10:**
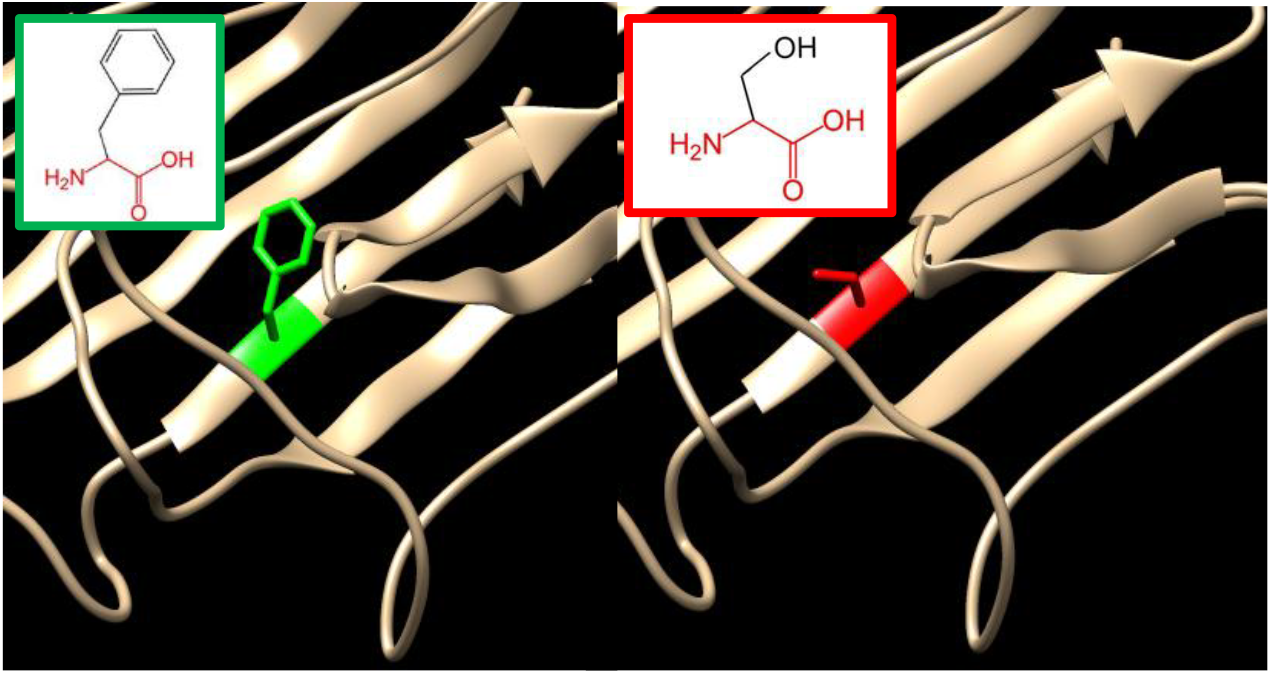
(F256S) The amino acid Phenylalanine change to Serine at 256.

**Figure 11:**
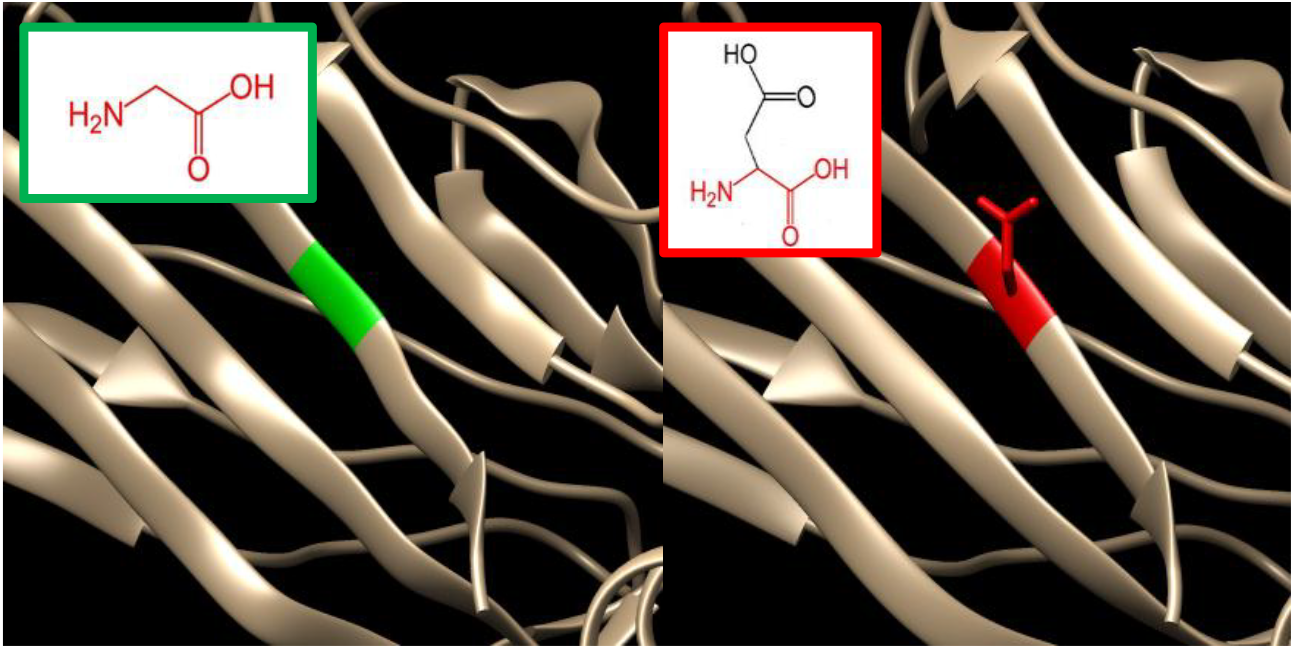
(G257D) The amino acid Glycine change to Aspartic Acid at 256.

GeneMANIA revealed that *CD40LG* has many vital functions: adaptive immune response based on somatic recombination of immune receptors built from immunoglobulin superfamily domains, B cell activation, B cell activation involved in immune response, B cell proliferation, cell activation involved in immune response, cell-type specific apoptotic process, endothelial cell apoptotic process, inflammatory response, isotype switching, leukocyte activation involved in immune response, leukocyte mediated immunity, lymphocyte activation involved in immune response, lymphocyte proliferation, mononuclear cell proliferation, positive regulation of apoptotic process, positive regulation of cell death, positive regulation of cytokine production, positive regulation of programmed cell death, production of molecular mediator of immune response, regulation of endothelial cell apoptotic process, somatic diversification of immunoglobulins involved in immune response, somatic recombination of immunoglobulin gene segments, somatic recombination of immunoglobulin genes involved in immune response, tumor necrosis factor receptor superfamily binding. We also analysed the correlations between significantly regulated genes, share similar protein domain, or participate to achieve similar function were illustrated by GeneMANIA and shown in figure 3, Tables (4&5).

**Table (4):** The *CD40LG* gene functions and its appearance in network and genome:

| Function | FDR | Genes in network | Genes in genome |
| --- | --- | --- | --- |
| necrotic cell death | 3.46E-05 | 4 | 22 |
| necroptotic process | 3.46E-05 | 4 | 18 |
| programmed necrotic cell death | 3.46E-05 | 4 | 19 |
| positive regulation of I-kappaB kinase/NF-kappaB signaling | 3.46E-05 | 6 | 138 |
| regulation of I-kappaB kinase/NF-kappaB signaling | 0.000103 | 6 | 185 |
| regulation of immune effector process | 0.000103 | 6 | 184 |
| I-kappaB kinase/NF-kappaB signaling | 0.000133 | 6 | 198 |
| positive regulation of cytokine production | 0.000151 | 6 | 207 |
| toll-like receptor 4 signaling pathway | 0.000183 | 5 | 107 |
| positive regulation of defense response | 0.000232 | 6 | 231 |
| cellular response to interleukin-4 | 0.000285 | 3 | 11 |
| CD40 receptor complex | 0.000285 | 3 | 11 |
| response to interleukin-4 | 0.00035 | 3 | 12 |
| toll-like receptor signaling pathway | 0.000405 | 5 | 137 |
| pattern recognition receptor signaling pathway | 0.000767 | 5 | 158 |
| innate immune response-activating signal transduction | 0.000767 | 5 | 160 |
| production of molecular mediator of immune response | 0.000879 | 4 | 69 |
| activation of innate immune response | 0.000879 | 5 | 168 |
| TRIF-dependent toll-like receptor signaling pathway | 0.00123 | 4 | 76 |
| MyD88-independent toll-like receptor signaling pathway | 0.001365 | 4 | 79 |
| regulation of endothelial cell apoptotic process | 0.001381 | 3 | 22 |
| cell-type specific apoptotic process | 0.001381 | 5 | 194 |
| positive regulation of innate immune response | 0.001381 | 5 | 191 |
| toll-like receptor 3 signaling pathway | 0.001387 | 4 | 83 |
| endothelial cell apoptotic process | 0.001887 | 3 | 25 |
| regulation of innate immune response | 0.003909 | 5 | 246 |
| B cell activation | 0.005198 | 4 | 119 |
| cytokine production involved in immune response | 0.006123 | 3 | 38 |
| regulation of production of molecular mediator of immune response | 0.009243 | 3 | 44 |
| regulation of cysteine-type endopeptidase activity | 0.015054 | 4 | 160 |
| side of membrane | 0.015581 | 4 | 163 |
| B cell proliferation | 0.015581 | 3 | 54 |
| regulation of defense response to virus | 0.02744 | 3 | 66 |
| response to interleukin-15 | 0.02744 | 2 | 10 |
| regulation of necrotic cell death | 0.02744 | 2 | 10 |
| response to virus | 0.032144 | 4 | 205 |
| lymphocyte activation involved in immune response | 0.032144 | 3 | 72 |
| cytoplasmic pattern recognition receptor signaling pathway in response to virus | 0.035169 | 2 | 12 |
| receptor complex | 0.035169 | 4 | 213 |
| cellular response to virus | 0.035169 | 2 | 12 |
| regulation of multi-organism process | 0.035534 | 4 | 216 |
| regulation of response to biotic stimulus | 0.049587 | 3 | 87 |
| positive regulation of apoptotic process | 0.058691 | 4 | 249 |
| regulation of endopeptidase activity | 0.059151 | 4 | 251 |
| leukocyte activation involved in immune response | 0.059317 | 3 | 96 |
| adaptive immune response based on somatic recombination of immune receptors built from immunoglobulin superfamily domains | 0.059317 | 3 | 98 |
| cell activation involved in immune response | 0.059317 | 3 | 96 |
| regulation of peptidase activity | 0.059317 | 4 | 258 |
| positive regulation of programmed cell death | 0.059317 | 4 | 254 |
| receptor signaling protein activity | 0.059317 | 3 | 98 |
| positive regulation of cell death | 0.065646 | 4 | 268 |
| T cell apoptotic process | 0.072358 | 2 | 20 |
| isotype switching | 0.072358 | 2 | 20 |
| somatic recombination of immunoglobulin genes involved in immune response | 0.072358 | 2 | 20 |
| somatic diversification of immunoglobulins involved in immune response | 0.072358 | 2 | 20 |
| inflammatory response | 0.072358 | 4 | 283 |
| regulation of leukocyte differentiation | 0.072358 | 3 | 109 |
| negative regulation of cytokine production | 0.07391 | 3 | 111 |
| tumor necrosis factor receptor superfamily binding | 0.074167 | 2 | 21 |
| positive regulation of NF-kappaB transcription factor activity | 0.074167 | 3 | 112 |
| defense response to virus | 0.080025 | 3 | 116 |
| NIK/NF-kappaB signaling | 0.083661 | 2 | 23 |
| B cell activation involved in immune response | 0.083661 | 2 | 23 |
| somatic recombination of immunoglobulin gene segments | 0.083661 | 2 | 23 |
| lymphocyte proliferation | 0.089235 | 3 | 123 |
| leukocyte mediated immunity | 0.09 | 3 | 124 |
| mononuclear cell proliferation | 0.090774 | 3 | 125 |

**\*FDR:** false discovery rate is greater than or equal to the probability that this is a false positive.

**Table (5):** The gene co-expressed, share domain and Interaction with *CD40LG* gene network:

**\*FDR:** false discovery rate is greater than or equal to the probability that this is a false positive. **Table (5) The gene co-expressed, share domain and Interaction with *CD40LG* gene network:**
| Gene 1 | Gene 2 | Weight | Network group |
| --- | --- | --- | --- |
| IL2RG | FASLG | 0.014963 | Co-expression |
| KCNA3 | IL4 | 0.013943 | Co-expression |
| NFKB2 | BIRC3 | 0.011734 | Co-expression |
| IL2RG | BIRC3 | 0.011704 | Co-expression |
| IL4 | CD40LG | 0.009129 | Co-expression |
| REL | IL4R | 0.011926 | Co-expression |
| KCNA3 | CD40LG | 0.010907 | Co-expression |
| ICOS | CD40 | 0.012159 | Co-expression |
| TRAF3 | ICOS | 0.008684 | Co-expression |
| FASLG | ICOS | 0.013282 | Co-expression |
| BIRC3 | ICOS | 0.008743 | Co-expression |
| BIRC3 | TRAF3 | 0.011122 | Co-expression |
| IL2RG | CD40 | 0.006743 | Co-expression |
| IL2RG | ICOS | 0.00355 | Co-expression |
| IL2RG | IL4R | 0.006168 | Co-expression |
| IL2RG | BIRC3 | 0.003711 | Co-expression |
| IL2RG | NFKB2 | 0.006361 | Co-expression |
| KCNA3 | IL2RG | 0.005632 | Co-expression |
| FASLG | ICOS | 0.021807 | Co-expression |
| TNF | JAK3 | 0.006956 | Co-expression |
| BIRC3 | CD40 | 0.017045 | Co-expression |
| NFKB2 | BIRC3 | 0.022512 | Co-expression |
| IL2RG | IL4R | 0.012735 | Co-expression |
| IL2RG | BIRC3 | 0.011602 | Co-expression |
| IL2RG | NFKB2 | 0.02143 | Co-expression |
| KCNA3 | CD40 | 0.017026 | Co-expression |
| BIRC3 | CD40 | 0.016166 | Co-expression |
| NFKB2 | CD40 | 0.019693 | Co-expression |
| IL2RG | CD40 | 0.008382 | Co-expression |
| IL2RG | ICOS | 0.007516 | Co-expression |
| IL2RG | BIRC3 | 0.005774 | Co-expression |
| IL2RG | NFKB2 | 0.006811 | Co-expression |
| KCNA3 | ICOS | 0.02028 | Co-expression |
| KCNA3 | IL2RG | 0.006984 | Co-expression |
| BIRC3 | CD40LG | 0.01894 | Co-expression |
| IL2RG | JAK3 | 0.008937 | Co-expression |
| BIRC3 | CD40 | 0.00956 | Co-expression |
| BIRC3 | TNF | 0.021513 | Co-expression |
| NFKB2 | CD40 | 0.008004 | Co-expression |
| NFKB2 | TNF | 0.012938 | Co-expression |
| NFKB2 | BIRC3 | 0.015358 | Co-expression |
| IL2RG | NFKB2 | 0.020683 | Co-expression |
| KCNA3 | MAP3K1 | 0.008914 | Co-expression |
| BIRC2 | TNF | 0.01865 | Co-expression |
| BIRC2 | BIRC3 | 0.028052 | Co-expression |
| IL4R | TRAF3 | 0.014878 | Co-expression |
| TNF | FASLG | 0.013372 | Co-expression |
| REL | JUND | 0.008833 | Co-expression |
| BIRC3 | TRAF3 | 0.013216 | Co-expression |
| NFKB2 | TRAF3 | 0.022174 | Co-expression |
| NFKB2 | TNF | 0.022091 | Co-expression |
| NFKB2 | BIRC3 | 0.013708 | Co-expression |
| MAP3K1 | REL | 0.01427 | Co-expression |
| IL4R | CD40 | 0.005425 | Co-expression |
| REL | IL4R | 0.011723 | Co-expression |
| BIRC3 | CD40 | 0.004387 | Co-expression |
| NFKB2 | CD40 | 0.011266 | Co-expression |
| IL2RG | CD40 | 0.005195 | Co-expression |
| IL2RG | IL4R | 0.007494 | Co-expression |
| BIRC2 | BIRC3 | 0.006445 | Co-expression |
| TNF | ICOS | 0.014079 | Co-expression |
| JUND | IL4R | 0.009701 | Co-expression |
| BIRC3 | CD40 | 0.010649 | Co-expression |
| RNF128 | TDP2 | 0.006606 | Co-expression |
| FASLG | CD40LG | 0.017517 | Co-expression |
| JAK3 | CD40LG | 0.020611 | Co-expression |
| TNF | CD40LG | 0.018848 | Co-expression |
| BIRC3 | CD40 | 0.019717 | Co-expression |
| NFKB2 | BIRC3 | 0.018493 | Co-expression |
| IL2RG | CD40 | 0.007105 | Co-expression |
| IL2RG | ICOS | 0.004861 | Co-expression |
| IL2RG | BIRC3 | 0.005204 | Co-expression |
| IL2RG | NFKB2 | 0.007049 | Co-expression |
| KCNA3 | CD40LG | 0.01689 | Co-expression |
| KCNA3 | ICOS | 0.016696 | Co-expression |
| KCNA3 | IL2RG | 0.006413 | Co-expression |
| NFKB2 | CD40 | 0.011105 | Co-expression |
| NFKB2 | IL4R | 0.01556 | Co-expression |
| NFKB2 | TNF | 0.011411 | Co-expression |
| NFKB2 | BIRC3 | 0.011462 | Co-expression |
| KCNA3 | BIRC3 | 0.006626 | Co-expression |
| BIRC2 | BIRC3 | 0.012962 | Co-expression |
| BIRC2 | NFKB2 | 0.00852 | Co-expression |
| BIRC2 | BIRC3 | 0.016683 | Co-expression |
| REL | TNF | 0.010127 | Co-expression |
| BIRC3 | CD40 | 0.00818 | Co-expression |
| IL2RG | JUND | 0.009352 | Co-expression |
| IL2RG | NFKB2 | 0.009404 | Co-expression |
| BIRC2 | CD40 | 0.010793 | Co-expression |
| BIRC2 | IL4 | 0.008467 | Co-expression |
| BIRC2 | TNF | 0.006199 | Co-expression |
| BIRC2 | BIRC3 | 0.007214 | Co-expression |
| BIRC3 | ICOS | 0.020492 | Co-expression |
| BIRC2 | REL | 0.010786 | Co-expression |
| REL | JUND | 0.023229 | Co-expression |
| IL2RG | REL | 0.012688 | Co-expression |
| BIRC2 | BIRC3 | 0.008191 | Co-expression |
| TNF | JAK3 | 0.026589 | Co-expression |
| BIRC3 | REL | 0.009161 | Co-expression |
| IL2RG | JAK3 | 0.011127 | Co-expression |
| IL2RG | IL4R | 0.004268 | Co-expression |
| IL2RG | TNF | 0.010053 | Co-expression |
| NFKB2 | CD40 | 0.005998 | Co-expression |
| NFKB2 | BIRC3 | 0.014023 | Co-expression |
| KCNA3 | MAP3K1 | 0.003699 | Co-expression |
| BIRC2 | TNF | 0.008398 | Co-expression |
| BIRC2 | BIRC3 | 0.015018 | Co-expression |
| FASLG | CD40LG | 0.015669 | Co-localization |
| JAK3 | CD40LG | 0.014456 | Co-localization |
| JAK3 | FASLG | 0.014457 | Co-localization |
| BIRC3 | CD40LG | 0.014902 | Co-localization |
| BIRC3 | FASLG | 0.020128 | Co-localization |
| KCNA3 | CD40LG | 0.013906 | Co-localization |
| KCNA3 | JAK3 | 0.021973 | Co-localization |
| JAK3 | CD40LG | 0.009828 | Co-localization |
| TNF | IL4R | 0.021817 | Co-localization |
| IL2RG | CD40LG | 0.004545 | Co-localization |
| KCNA3 | CD40LG | 0.008507 | Co-localization |
| KCNA3 | IL2RG | 0.005577 | Co-localization |
| BIRC2 | BIRC3 | 0.015573 | Co-localization |
| CD40 | CD40LG | 0.192562 | Pathway |
| ICOS | CD40LG | 0.129156 | Pathway |
| TRAF3 | CD40LG | 0.060989 | Pathway |
| TRAF3 | CD40 | 0.129558 | Pathway |
| IL4 | CD40LG | 0.05932 | Pathway |
| JAK3 | CD40LG | 0.023855 | Pathway |
| JAK3 | IL4 | 0.01561 | Pathway |
| IL4R | CD40LG | 0.049297 | Pathway |
| IL4R | IL4 | 0.03226 | Pathway |
| IL4R | JAK3 | 0.012973 | Pathway |
| TNF | FASLG | 0.01287 | Pathway |
| JUND | CD40LG | 0.055153 | Pathway |
| REL | CD40LG | 0.05313 | Pathway |
| MAP4K4 | CD40LG | 0.048895 | Pathway |
| MAP4K4 | CD40 | 0.103865 | Pathway |
| MAP4K4 | TRAF3 | 0.032897 | Pathway |
| MAP4K4 | TNF | 0.012563 | Pathway |
| BIRC3 | FASLG | 0.023192 | Pathway |
| BIRC3 | TNF | 0.010785 | Pathway |
| BIRC3 | MAP4K4 | 0.022639 | Pathway |
| NFKB2 | CD40LG | 0.042237 | Pathway |
| NFKB2 | REL | 0.024756 | Pathway |
| MAP3K1 | CD40LG | 0.028289 | Pathway |
| MAP3K1 | CD40 | 0.060093 | Pathway |
| MAP3K1 | TRAF3 | 0.019033 | Pathway |
| MAP3K1 | TNF | 0.007269 | Pathway |
| MAP3K1 | MAP4K4 | 0.015259 | Pathway |
| IL2RG | CD40LG | 0.028033 | Pathway |
| IL2RG | IL4 | 0.018345 | Pathway |
| IL2RG | JAK3 | 0.007377 | Pathway |
| IL2RG | IL4R | 0.015245 | Pathway |
| BIRC2 | TNF | 0.016131 | Pathway |
| BIRC2 | BIRC3 | 0.029068 | Pathway |
| CD40 | CD40LG | 0.046437 | Pathway |
| TRAF3 | CD40LG | 0.119048 | Pathway |
| TRAF3 | CD40 | 0.151661 | Pathway |
| IL4 | CD40LG | 0.026394 | Pathway |
| JAK3 | CD40LG | 0.024033 | Pathway |
| JAK3 | CD40 | 0.030616 | Pathway |
| JAK3 | IL4 | 0.017402 | Pathway |
| IL4R | CD40LG | 0.041845 | Pathway |
| IL4R | IL4 | 0.0303 | Pathway |
| IL4R | JAK3 | 0.027589 | Pathway |
| TDP2 | CD40LG | 0.136808 | Pathway |
| TDP2 | CD40 | 0.174285 | Pathway |
| MAP4K4 | TNF | 0.041482 | Pathway |
| BIRC3 | CD40LG | 0.032145 | Pathway |
| BIRC3 | CD40 | 0.040951 | Pathway |
| BIRC3 | TRAF3 | 0.104984 | Pathway |
| BIRC3 | TNF | 0.016886 | Pathway |
| BIRC3 | MAP4K4 | 0.069639 | Pathway |
| MAP3K1 | CD40LG | 0.031848 | Pathway |
| MAP3K1 | CD40 | 0.040573 | Pathway |
| IL2RG | CD40LG | 0.019385 | Pathway |
| IL2RG | IL4 | 0.014036 | Pathway |
| IL2RG | JAK3 | 0.012781 | Pathway |
| IL2RG | IL4R | 0.022253 | Pathway |
| RNF128 | IL4 | 0.115976 | Pathway |
| BIRC2 | CD40LG | 0.076342 | Pathway |
| BIRC2 | CD40 | 0.097255 | Pathway |
| BIRC2 | TRAF3 | 0.249331 | Pathway |
| BIRC2 | TNF | 0.040103 | Pathway |
| BIRC2 | BIRC3 | 0.067323 | Pathway |
| CD40 | CD40LG | 0.607524 | Physical Interactions |
| TRAF3 | CD40 | 0.095902 | Physical Interactions |
| IL4R | IL4 | 0.796225 | Physical Interactions |
| BIRC3 | CD40 | 0.330114 | Physical Interactions |
| NFKB2 | REL | 0.047741 | Physical Interactions |
| IL2RG | JAK3 | 0.53926 | Physical Interactions |
| CD40 | CD40LG | 0.08088 | Physical Interactions |
| TRAF3 | CD40 | 0.013307 | Physical Interactions |
| HPR | CD40LG | 0.523536 | Physical Interactions |
| IL4R | CD40 | 0.054789 | Physical Interactions |
| IL4R | IL4 | 0.343558 | Physical Interactions |
| IL4R | JAK3 | 0.050891 | Physical Interactions |
| TDP2 | CD40 | 0.07344 | Physical Interactions |
| TDP2 | TRAF3 | 0.025738 | Physical Interactions |
| BIRC3 | CD40LG | 0.039862 | Physical Interactions |
| BIRC3 | CD40 | 0.018714 | Physical Interactions |
| BIRC3 | TRAF3 | 0.006558 | Physical Interactions |
| BIRC3 | REL | 0.020398 | Physical Interactions |
| NFKB2 | REL | 0.048602 | Physical Interactions |
| MAP3K1 | MAP4K4 | 0.019982 | Physical Interactions |
| IL2RG | JAK3 | 0.04765 | Physical Interactions |
| IL2RG | IL4R | 0.074022 | Physical Interactions |
| RNF128 | CD40LG | 0.164195 | Physical Interactions |
| BIRC2 | CD40LG | 0.032488 | Physical Interactions |
| BIRC2 | CD40 | 0.015252 | Physical Interactions |
| BIRC2 | TRAF3 | 0.005345 | Physical Interactions |
| BIRC2 | TNF | 0.010899 | Physical Interactions |
| BIRC2 | BIRC3 | 0.007517 | Physical Interactions |
| BIRC2 | NFKB2 | 0.017911 | Physical Interactions |
| TRAF3 | CD40 | 0.475875 | Physical Interactions |
| IL4R | IL4 | 0.372073 | Physical Interactions |
| IL2RG | IL4 | 0.399409 | Physical Interactions |
| IL4R | IL4 | 1 | Physical Interactions |
| TNF | ICOS | 0.102404 | Physical Interactions |
| NFKB2 | REL | 0.173546 | Physical Interactions |
| TRAF3 | CD40 | 0.036334 | Physical Interactions |
| IL4R | IL4 | 0.454577 | Physical Interactions |
| NFKB2 | REL | 0.00236 | Physical Interactions |
| CD40 | CD40LG | 0.105262 | Physical Interactions |
| TRAF3 | CD40 | 0.023258 | Physical Interactions |
| JAK3 | CD40 | 0.033453 | Physical Interactions |
| IL4R | CD40 | 0.04218 | Physical Interactions |
| IL4R | IL4 | 0.104439 | Physical Interactions |
| IL4R | JAK3 | 0.035954 | Physical Interactions |
| TDP2 | CD40 | 0.077995 | Physical Interactions |
| TDP2 | TRAF3 | 0.046223 | Physical Interactions |
| BIRC3 | CD40 | 0.078458 | Physical Interactions |
| NFKB2 | REL | 0.037047 | Physical Interactions |
| IL2RG | IL4 | 0.095945 | Physical Interactions |
| IL2RG | JAK3 | 0.03303 | Physical Interactions |
| IL2RG | IL4R | 0.041646 | Physical Interactions |
| RNF128 | CD40LG | 0.306793 | Physical Interactions |
| BIRC2 | CD40 | 0.041099 | Physical Interactions |
| TRAF3 | CD40 | 0.101777 | Predicted |
| NFKB2 | REL | 0.060585 | Predicted |
| BIRC2 | BIRC3 | 0.482262 | Predicted |
| FASLG | CD40LG | 0.245935 | Predicted |
| FASLG | CD40 | 0.03527 | Predicted |
| JAK3 | CD40 | 0.046533 | Predicted |
| IL4R | CD40 | 0.188654 | Predicted |
| TNF | CD40LG | 0.152204 | Predicted |
| TNF | CD40 | 0.021828 | Predicted |
| BIRC3 | TRAF3 | 0.068934 | Predicted |
| NFKB2 | TRAF3 | 0.024427 | Predicted |
| BIRC2 | CD40 | 0.048271 | Predicted |
| IL2RG | IL4 | 0.664124 | Predicted |
| MAP3K1 | MAP4K4 | 0.258003 | Predicted |
| IL2RG | JAK3 | 1 | Predicted |
| FASLG | CD40LG | 0.041645 | Shared protein domains |
| TNF | CD40LG | 0.038418 | Shared protein domains |
| TNF | FASLG | 0.050344 | Shared protein domains |
| NFKB2 | REL | 0.049596 | Shared protein domains |
| IL2RG | IL4R | 0.041143 | Shared protein domains |
| BIRC2 | BIRC3 | 0.047106 | Shared protein domains |
| FASLG | CD40LG | 0.064011 | Shared protein domains |
| TNF | CD40LG | 0.064011 | Shared protein domains |
| TNF | FASLG | 0.064011 | Shared protein domains |
| NFKB2 | REL | 0.049118 | Shared protein domains |
| MAP3K1 | MAP4K4 | 0.003437 | Shared protein domains |
| BIRC2 | BIRC3 | 0.056869 | Shared protein domains |

We also used ConSurf web server; the nsSNPs that are located at highly conserved amino acid positions tend to be more deleterious than nsSNPs that are located at non-conserved sites. (Figure2).

Some study revealed that, patients with Hyper IgM syndrome are susceptible to malignancies of neuroendocrine origin know as neuroendocrine tumors (NETs), of all primary immune deficiency diseases, NETs appeared to be unique to XHIGM patients[35] The mechanism behind this association remains unclear, so we hope that this study will help in understanding and diagnosis of this disorder. We also believe that, any mutations in *CD40LG* may be a part of molecular events involved in the severity of Sickle cell disease and Down syndrome. [36, 56]

This is the first in silico analysis of *CD40LG* gene that used functional analysis to determine the deleterious SNPs. Our study matches the result from the NCBI that 6 SNPs are pathogenic (W140G and G227V are pathogenic while G144E, L155P, Y169C and F256S likely pathogenic) yet the remaining 2 SNPs (Y170C and G257D) were untested in the NCBI but found deleterious in our study. These SNPs may now be used as diagnostic markers for hyper IgM syndrome and may help in the understanding of the associated diseases. Finally some gratitude of wet lab techniques is suggested to support our findings.

## Conclusion

In this study the influence of functional SNPs in the *CD40LG* gene was thoroughly investigated through different bioinformatics prediction softwares. A total of 8 novel mutations (W140G, G144E, L155P, Y169C, Y170C, G227V, F256S, and G257D) were predicted to be responsible for the structural and functional modifications of *CD40LG* protein. It is evident from multiple in silico analysis tools that; these 8 nsSNPs might serve as, a novel diagnostic markers of Hyper-IgM syndrome. And can assist in pharmacogenomics by identifying high risk SNP variants contributing to drug response as well as developing novel therapeutic elements. And could help in the overall understanding of this disease.

## Acknowledgment

The authors wish to acknowledgment the enthusiastic cooperation of Africa City of Technology - Sudan.

## References

[1] J. Johnson, A. H. Filipovich, and K. Zhang, “X-Linked Hyper IgM Syndrome,” in *GeneReviews((R))*, M. P. Adam, H. H. Ardinger, R. A. Pagon, S. E. Wallace, L. J. H. Bean, K. Stephens, et al., Eds., ed Seattle (WA): University of Washington, Seattle University of Washington, Seattle. GeneReviews is a registered trademark of the University of Washington, Seattle. All rights reserved., 1993.

[2] L. C. Schneider, “X-linked hyper IgM syndrome,” Clin Rev Allergy Immunol, vol. 19, pp. 205–15, Oct 2000.

[3] D. Buchbinder, S. Park, and D. Nugent, “X-linked hyper IgM syndrome: a novel sequence variant associated with an atypical mild phenotype,” J Pediatr Hematol Oncol, vol. 34, pp. e212–4, Jul 2012.

[4] M. Groeneweg, A. C. Lankester, and R. G. Bredius, “[From gene to disease; CD40 ligand deficiency as the cause of X-linked hyper-IgM-syndrome],” Ned Tijdschr Geneeskd, vol. 147, pp. 1009-11, May 24 2003.

[5] R. L. Fuleihan, “Hyper IgM syndrome: the other side of the coin,” Curr Opin Pediatr, vol. 13, pp. 528–32, Dec 2001.

[6] R. L. Fuleihan, “The hyper IgM syndrome,” Curr Allergy Asthma Rep, vol. 1, pp. 445–50, Sep 2001.

[7] S. Rigaud, E. Lopez-Granados, S. Siberil, G. Gloire, N. Lambert, C. Lenoir, et al., “Human X-linked variable immunodeficiency caused by a hypomorphic mutation in XIAP in association with a rare polymorphism in CD40LG,” Blood, vol. 118, pp. 252–61, Jul 14 2011.

[8] A. Rawat, B. Mathew, V. Pandiarajan, A. Jindal, M. Sharma, D. Suri, et al., “Clinical and molecular features of X-linked hyper IgM syndrome - An experience from North India,” Clin Immunol, vol. 195, pp. 59–66, Oct 2018.

[9] C. Aloui, A. Prigent, S. Tariket, C. Sut, J. Fagan, F. Cognasse, et al., “Levels of human platelet-derived soluble CD40 ligand depend on haplotypes of CD40LG-CD40-ITGA2,” Sci Rep, vol. 6, p. 24715, Apr 20 2016.

[10] D. Nandan, V. K. Nag, N. Trivedi, and S. Singh, “X-linked Hyper-IgM Syndrome with Bronchiectasis,” J Lab Physicians, vol. 6, pp. 114–6, Jul 2014.

[11] K. Y. Qiu, X. Y. Liao, R. H. Wu, K. Huang, J. P. Fang, and D. H. Zhou, “X-linked Hyper-IgM Syndrome: A Phenotype of Crohn’s Disease with Hemophagocytic Lymphohistiocytosis,” Pediatr Hematol Oncol, vol. 34, pp. 428–434, Nov 2017.

[12] A. Jasinska, K. Kalwak, J. Trelinska, M. Borowiec, B. Piatosa, K. Zeman, et al., “Successful haploidentical PBSCT with subsequent T-cell addbacks in a boy with HyperIgM syndrome presenting as severe congenital neutropenia,” Pediatr Transplant, vol. 17, pp. E37–40, Feb 2013.

[13] P. Pacharn, W. Phongsamart, B. Boonyawat, O. Jirapongsananuruk, N. Visitsunthorn, and K. Chokephaibulkit, “Disseminated cryptococcosis in two boys with novel mutation of CD40 Ligand-Associated X-linked hyper-IgM syndrome,” Asian Pac J Allergy Immunol, Oct 15 2018.

[14] H. Y. Tsai, H. H. Yu, Y. H. Chien, K. H. Chu, Y. L. Lau, J. H. Lee, et al., “X-linked hyper-IgM syndrome with CD40LG mutation: two case reports and literature review in Taiwanese patients,” J Microbiol Immunol Infect, vol. 48, pp. 113–8, Feb 2015.

[15] E. Yousef and M. Arshad Alvi, “Hyper IgM Syndrome with low IgM and thrombocytosis: an unusual case of immunodeficiency,” Eur Ann Allergy Clin Immunol, vol. 48, pp. 194–6, Sep 2016.

[16] C. Aloui, C. Sut, F. Cognasse, V. Granados, M. Hassine, T. Chakroun, et al., “Development of a highly resolutive method, using a double quadruplex tetra-primer-ARMS-PCR coupled with capillary electrophoresis to study CD40LG polymorphisms,” Mol Cell Probes, vol. 29, pp. 335–342, Dec 2015.

[17] C. Aloui, C. Sut, A. Prigent, J. Fagan, F. Cognasse, V. Granados-Herbepin, et al., “Are polymorphisms of the immunoregulatory factor CD40LG implicated in acute transfusion reactions?,” Sci Rep, vol. 4, p. 7239, Nov 28 2014.

[18] M. P. Cicalese, J. Gerosa, M. Baronio, D. Montin, F. Licciardi, A. Soresina, et al., “Circulating Follicular Helper and Follicular Regulatory T Cells Are Severely Compromised in Human CD40 Deficiency: A Case Report,” Front Immunol, vol. 9, p. 1761, 2018.

[19] N. Hubbard, D. Hagin, K. Sommer, Y. Song, I. Khan, C. Clough, et al., “Targeted gene editing restores regulated CD40L function in X-linked hyper-IgM syndrome,” Blood, vol. 127, pp. 2513-22, May 26 2016.

[20] K. Imai, M. Shimadzu, T. Kubota, T. Morio, T. Matsunaga, Y. D. Park, et al., “Female hyper IgM syndrome type 1 with a chromosomal translocation disrupting CD40LG,” Biochim Biophys Acta, vol. 1762, pp. 335–40, Mar 2006.

[21] A. Horrillo, T. Fontela, E. G. Arias-Salgado, D. Llobat, G. Porras, M. S. Ayuso, et al., “Generation of mice with conditional ablation of the Cd40lg gene: new insights on the role of CD40L,” Transgenic Res, vol. 23, pp. 53–66, Feb 2014.

[22] P. A. Apoil, E. Kuhlein, A. Robert, H. Rubie, and A. Blancher, “HIGM syndrome caused by insertion of an AluYb8 element in exon 1 of the CD40LG gene,” Immunogenetics, vol. 59, pp. 17–23, Jan 2007.

[23] L. Martinez-Martinez, C. Gonzalez-Santesteban, I. Badell, and O. de la Calle-Martin, “The polymorphism p.G219R of CD40L does not cause immunological alterations in vivo: conclusions from a X-linked hyper IgM syndrome kindred,” Mol Immunol, vol. 52, pp. 237–41, Oct 2012.

[24] Q. Dong, J. Zhao, Z. Yao, X. Liu, and H. He, “A Case Report of X-Linked Hyperimmunoglobulin M Syndrome with Lipoma Arborescens of Knees,” Case Rep Med, vol. 2016, p. 5797232, 2016.

[25] T. T. Franca, L. F. B. Leite, T. A. Maximo, C. G. Lambert, N. B. Zurro, W. C. N. Forte, et al., “A Novel de Novo Mutation in the CD40 Ligand Gene in a Patient With a Mild X-Linked Hyper-IgM Phenotype Initially Diagnosed as CVID: New Aspects of Old Diseases,” Front Pediatr, vol. 6, p. 130, 2018.

[26] A. Hewagama, G. Gorelik, D. Patel, P. Liyanarachchi, W. J. McCune, E. Somers, et al., “Overexpression of X-linked genes in T cells from women with lupus,” J Autoimmun, vol. 41, pp. 60–71, Mar 2013.

[27] H. Y. Kim, T. M. Um, and H. J. Park, “A Novel Mutation in CD40LG Gene Causing X-Linked Hyper IgM Syndrome,” Indian J Pediatr, vol. 85, pp. 788–789, Sep 2018.

[28] H. Ouadani, I. Ben-Mustapha, M. Ben-ali, L. Ben-khemis, B. Largueche, R. Boussoffara, et al., “Novel and recurrent AID mutations underlie prevalent autosomal recessive form of HIGM in consanguineous patients,” Immunogenetics, vol. 68, pp. 19–28, Jan 2016.

[29] A. Rangel-Santos, V. L. Wakim, C. M. Jacob, A. C. Pastorino, J. M. Cunha, A. C. Collanieri, et al., “Molecular characterization of patients with X-linked Hyper-IgM syndrome: description of two novel CD40L mutations,” Scand J Immunol, vol. 69, pp. 169–73, Feb 2009.

[30] L. L. Wang, W. Zhou, W. Zhao, Z. Q. Tian, W. F. Wang, X. F. Wang, et al., “Clinical features and genetic analysis of 20 Chinese patients with X-linked hyper-IgM syndrome,” J Immunol Res, vol. 2014, p. 683160, 2014.

[31] L. Yu, X. Wang, Y. Wang, and J. Wang, “Identification of two novel mutations in patients with X-linked primary immunodeficiencies,” Fetal Pediatr Pathol, vol. 34, pp. 91–8, Apr 2015.

[32] S. C. Lin, S. D. Shyur, W. I. Lee, Y. C. Ma, and L. H. Huang, “X-linked hyper-immunoglobulin M syndrome: molecular genetic study and long-time follow-up of three generations of a Chinese family,” Int Arch Allergy Immunol, vol. 140, pp. 1-8, 2006.

[33] N. Kutukculer, N. E. Karaca, G. Aksu, A. Aykut, E. Pariltay, and O. Cogulu, “An X-Linked Hyper-IgM Patient Followed Successfully for 23 Years without Hematopoietic Stem Cell Transplantation,” Case Reports Immunol, vol. 2018, p. 6897935, 2018.

[34] X. Liu, K. Zhou, D. Yu, X. Cai, Y. Hua, H. Zhou, et al., “A delayed diagnosis of X-linked hyper IgM syndrome complicated with toxoplasmic encephalitis in a child: A case report and literature review,” Medicine (Baltimore), vol. 96, p. e8989, Dec 2017.

[35] R. E. Nicolaides and M. T. de la Morena, “Inherited and acquired clinical phenotypes associated with neuroendocrine tumors,” Curr Opin Allergy Clin Immunol, vol. 17, pp. 431–442, Dec 2017.

[36] D. L. Zanette, R. P. Santiago, I. P. R. Leite, S. S. Santana, C. da Guarda, V. V. Maffili, et al., “Differential gene expression analysis of sickle cell anemia in steady and crisis state,” Ann Hum Genet, Jan 30 2019.

[37] B. M. Javierre and B. Richardson, “A new epigenetic challenge: systemic lupus erythematosus,” Adv Exp Med Biol, vol. 711, pp. 117–36, 2011.

[38] C. George Priya Doss, C. Sudandiradoss, R. Rajasekaran, P. Choudhury, P. Sinha, P. Hota, et al., “Applications of computational algorithm tools to identify functional SNPs,” Funct Integr Genomics, vol. 8, pp. 309–16, Nov 2008.

[39] X. Chen and J. T. Chang, “Planning bioinformatics workflows using an expert system,” Bioinformatics, vol. 33, pp. 1210-1215, Apr 15 2017.

[40] A. Bajard, S. Chabaud, C. Cornu, A. C. Castellan, S. Malik, P. Kurbatova, et al., “An in silico approach helped to identify the best experimental design, population, and outcome for future randomized clinical trials,” J Clin Epidemiol, vol. 69, pp. 125–36, Jan 2016.

[41] J. Wang, G. S. Pang, S. S. Chong, and C. G. Lee, “SNP web resources and their potential applications in personalized medicine,” Curr Drug Metab, vol. 13, pp. 978–90, Sep 1 2012.

[42] D. Gefel, I. Maslovsky, and J. Hillel, “[Application of single nucleotide polymorphisms (SNPs) for the detection of genes involved in the control of complex diseases],” Harefuah, vol. 147, pp. 449-54, 476, May 2008.

[43] N. L. Sim, P. Kumar, J. Hu, S. Henikoff, G. Schneider, and P. C. Ng, “SIFT web server: predicting effects of amino acid substitutions on proteins,” Nucleic Acids Res, vol. 40, pp. W452–7, Jul 2012.

[44] E. Capriotti and R. B. Altman, “Improving the prediction of disease-related variants using protein three-dimensional structure,” BMC Bioinformatics, vol. 12 Suppl 4, p. S3, 2011.

[45] Y. Choi, G. E. Sims, S. Murphy, J. R. Miller, and A. P. Chan, “Predicting the functional effect of amino acid substitutions and indels,” PLoS One, vol. 7, p. e46688, 2012.

[46] M. Hecht, Y. Bromberg, and B. Rost, “Better prediction of functional effects for sequence variants,” BMC Genomics, vol. 16 Suppl 8, p. S1, 2015.

[47] R. Calabrese, E. Capriotti, P. Fariselli, P. L. Martelli, and R. Casadio, “Functional annotations improve the predictive score of human disease-related mutations in proteins,” Hum Mutat, vol. 30, pp. 1237–44, Aug 2009.

[48] E. Capriotti, P. Fariselli, and R. Casadio, “I-Mutant2.0: predicting stability changes upon mutation from the protein sequence or structure,” Nucleic Acids Res, vol. 33, pp. W306–10, Jul 1 2005.

[49] J. Cheng, A. Randall, and P. Baldi, “Prediction of protein stability changes for single-site mutations using support vector machines,” Proteins, vol. 62, pp. 1125–32, Mar 1 2006.

[50] D. Warde-Farley, S. L. Donaldson, O. Comes, K. Zuberi, R. Badrawi, P. Chao, et al., “The GeneMANIA prediction server: biological network integration for gene prioritization and predicting gene function,” Nucleic Acids Res, vol. 38, pp. W214–20, Jul 2010.

[51] S. Wang, W. Li, S. Liu, and J. Xu, “RaptorX-Property: a web server for protein structure property prediction,” Nucleic Acids Res, vol. 44, pp. W430–5, Jul 8 2016.

[52] E. F. Pettersen, T. D. Goddard, C. C. Huang, G. S. Couch, D. M. Greenblatt, E. C. Meng, et al., “UCSF Chimera--a visualization system for exploratory research and analysis,” J Comput Chem, vol. 25, pp. 1605–12, Oct 2004.

[53] H. Ashkenazy, E. Erez, E. Martz, T. Pupko, and N. Ben-Tal, “ConSurf 2010: calculating evolutionary conservation in sequence and structure of proteins and nucleic acids,” Nucleic Acids Res, vol. 38, pp. W529–33, Jul 2010.

[54] H. Ashkenazy, S. Abadi, E. Martz, O. Chay, I. Mayrose, T. Pupko, et al., “ConSurf 2016: an improved methodology to estimate and visualize evolutionary conservation in macromolecules,” Nucleic Acids Res, vol. 44, pp. W344–50, Jul 8 2016.

[55] J. Jylhava, C. Eklund, M. Jylha, A. Hervonen, and M. Hurme, “Expression profiling of immune-associated genes in peripheral blood mononuclear cells reveals baseline differences in co-stimulatory signalling between nonagenarians and younger controls: the vitality 90+ study,” Biogerontology, vol. 11, pp. 671–7, Dec 2010.

[56] B. L. Zampieri, J. M. Biselli-Perico, J. E. de Souza, M. C. Burger, W. A. Silva Junior, E. M. Goloni-Bertollo, et al., “Altered expression of immune-related genes in children with Down syndrome,” PLoS One, vol. 9, p. e107218, 2014.

